# Afadin Sorts Different Retinal Neuron Types into Accurate Cellular Layers

**DOI:** 10.1101/2024.12.24.630272

**Authors:** Matthew R. Lum, Sachin H. Patel, Hannah K. Graham, Mengya Zhao, Yujuan Yi, Liang Li, Melissa Yao, Anna La Torre, Luca Della Santina, Ying Han, Yang Hu, Derek S. Welsbie, Xin Duan

## Abstract

Neurons use cell-adhesion molecules (CAMs) to interact with other neurons and the extracellular environment: the combination of CAMs specifies migration patterns, neuronal morphologies, and synaptic connections across diverse neuron types. Yet little is known regarding the intracellular signaling cascade mediating the CAM recognitions at the cell surface across different neuron types. In this study, we investigated the neural developmental role of Afadin^1–4^, a cytosolic adapter protein that connects multiple CAM families to intracellular F-actin. We introduced the conditional Afadin mutant^5^ to an embryonic retinal Cre, Six3-Cre^6–8^. We reported that the mutants lead to the scrambled retinal neuron distribution, including Bipolar Cells (BCs), Amacrine Cells (ACs), and retinal ganglion cells (RGCs), across three cellular layers of the retina. This scrambled pattern was first reported here at neuron-type resolution. Importantly, the mutants do not display deficits for BCs, ACs, or RGCs in terms of neural fate specifications or survival. Additionally, the displayed RGC types still maintain synaptic partners with putative AC types, indicating that other molecular determinants instruct synaptic choices independent of Afadin. Lastly, there is a significant decline in visual function and mis-targeting of RGC axons to incorrect zones of the superior colliculus, one of the major retinorecipient areas. Collectively, our study uncovers a unique cellular role of Afadin in sorting retinal neuron types into proper cellular layers as the structural basis for orderly visual processing.

## INTRODUCTION

The central nervous system (CNS) consists of complex circuits that are established during development and fine-tuned through both activity-dependent and apoptotic mechanisms. However, how neurons form these complex circuits—and, more specifically, how cell surface molecules promote correct neuronal migration and synapse formation—is not fully resolved. Cell adhesion molecules (CAMs) have been shown to be key mediators of CNS lamination and synaptogenesis. Work from ours and others focused on using the inner retina of the mouse as a developmental system to understand mechanisms regulating these developmental questions^9–13^. Specifically, our past studies showed that Type II Cadherin (Cdhs), in combination, play key roles in establishing appropriate synapses between retinal ganglion cells (RGCs) and bipolar cells (BCs), as well as RGCs with amacrine cells (ACs)^14,15^. Among the molecular machinery that composes the adherens junction (AJ) complex, β-catenin has been shown to be important in neuronal laminar organization, leading to embryonic deficits and major retinal neuron loss^16^.

While we assume that all intracellular components of cell-surface adhesion complexes are critical for retinal neuron survival and patterning, not all components of the AJ complex equally regulate the same aspects of these developmental programs. Afadin is a cytosolic adaptor protein that links Nectin, a Ca^2+^ independent immunoglobulin-like CAM, to F-actin microfilaments in the cytoskeleton^1–3,17^. Afadin recruits cadherins to AJs mediated by Nectin, p120-catenin, and α-catenin^2,3^. While Afadin has multiple direct interactions with AJ proteins, it is not a core part of the cadherin–catenin complex^18^.

Past genetic studies in the mouse CNS showed that loss of Afadin in both the hippocampal and cortical regions of the mouse brain leads to a decrease in dendritic spine density and number of synapses, with variable effects on dendritic arborization^5^. Additionally, Afadin plays an important role in cortical lamination: deletion of Afadin in the mouse telencephalon leads to cellular mislocalization and a resultant double-cortex^19,20^. Notably, the *drosophila* homologue of Afadin is called *Canoe*^21,22^, where the mutant phenotypes in the ommatidial eye were likely closely tied to the disruption of cellular junctions or synaptic complex, though given the broad role of Afadin (Canoe), they may also be due to other cell-surface signaling pathways^23^.

The mouse neural retina offers a laminarly organized structure and well-characterized cellular composition across three cellular layers. During development, retinal progenitor cells span the retinal neuroepithelium via basal and apical processes, proliferate via asymmetric and symmetric divisions at the ventricular surface, and differentiate into six neuronal types via both transcriptional regulatory networks and environmental cues—these include rod and cone photoreceptors (PR), horizontal cells (HC), bipolar cells (BC), amacrine cells (AC), retinal ganglion cells (RGC), and one glial cell type, Müller glia (MGs)^24–27^. Thus, the distinct locations and temporal order offer a clear system to examine the roles of multifaceted molecules in every step of development, such as that for Afadin. By restricting the roles of Afadin into restricted RGC subsets or AC subsets, our recent study linked Afadin to the combinatorial Cdh complex that enables the selective RGC-AC synaptic choice^15^. Yet it is unknown what role Afadin plays in neuronal migration, neuronal layer sorting, and brain target selection. Here, we utilized a developmental neural retina-specific Cre driver (Six3^Cre^)^6–8^ to generate a conditional Afadin mutant (Six3^Cre^; Afadin^F/F^). This conditional mutant allows us to characterize the role of Afadin in early development. Here, we report that the Afadin mutant significantly alters retinal neuronal migration and neuronal layer sorting, though it has little effect on cellular differentiation within the inner retina.

## RESULTS

### Early Afadin conditional mutant scramble retinal neuron layer organization

We utilized a murine conditional knockout model in which Exon 2 of Afadin was flanked by the LoxP sites (Figure 1U)^5^. Cre-mediated recombination results in the excision of exon 2, resulting in a frameshift mutation and a premature stop codon. These conditional alleles were crossed with Six3^Cre^ transgenic mice to mediate gene deletion, particularly within the developing retinal neuroepithelium starting at E9 ^6–8^. Upon conditional knockout, Afadin mutants (hereafter referred to as Afadin-cKO^Ret^) displayed aberrant lamination patterning at four different postnatal time points examined: P2, P7, P14, and P60 (Figure 1E-H). At P2, control retinas (Afadin^F/F^) exist as a singularly laminated piece of tissue resulting from the proliferation and early differentiation of early-born retinal neurons (Figure 1A). In contrast, Afadin-cKO^Ret^, the central retina is disrupted and contains rosette-like structures with regions devoid of cells at P2 (Figure 1E). At P7, the inner and outer plexiform layers are evident in control mice (Figure 1B) but notably disrupted in Afadin-cKO^Ret^, with the OPL omitted and instead existing instead as a singular fused nuclear layer (fused INL/ONL) (Figure 1F). P7 and older mutants also show columns of neurons (Figure 1F, arrow; Figure 2F) spanning the inner plexiform layer (IPL) and rosettes in the fused INL/ONL (Figure 1F, asterisks). By P14, most, if not all, retinal neuron types and Muller glial cells have been established^24,25^. To determine whether loss of Afadin affects retinal neuron densities or cell fate differentiation in addition to cellular layer organization, we examined mouse retinas and quantified neuronal subtypes in controls and mutants at P14 using well-established molecular markers: RBPMS to label RGCs, Chx10 to label BCs, and AP2α to label ACs. We found that the densities of the three major inner retinal neuronal types did not differ significantly between controls and mutants (Figure 1T), indicating retinal neurons in Afadin- cKO^Ret^ differentiate and proliferate via expected proportions, and Afadin is likely not involved in fate determination or cell survival regulations. When we used P14 mice to quantify the mis-localization of retinal types across three cellular layers, RGCs, ACs, and BCs were significantly scrambled across three cellular layers, compared to control (Figure 1S). Collectively, these results revealed the roles of Afadin in sorting inner retinal neurons into proper layers, likely integrating the positioning cues from the environment when specifying the layers.

**Figure 1.**
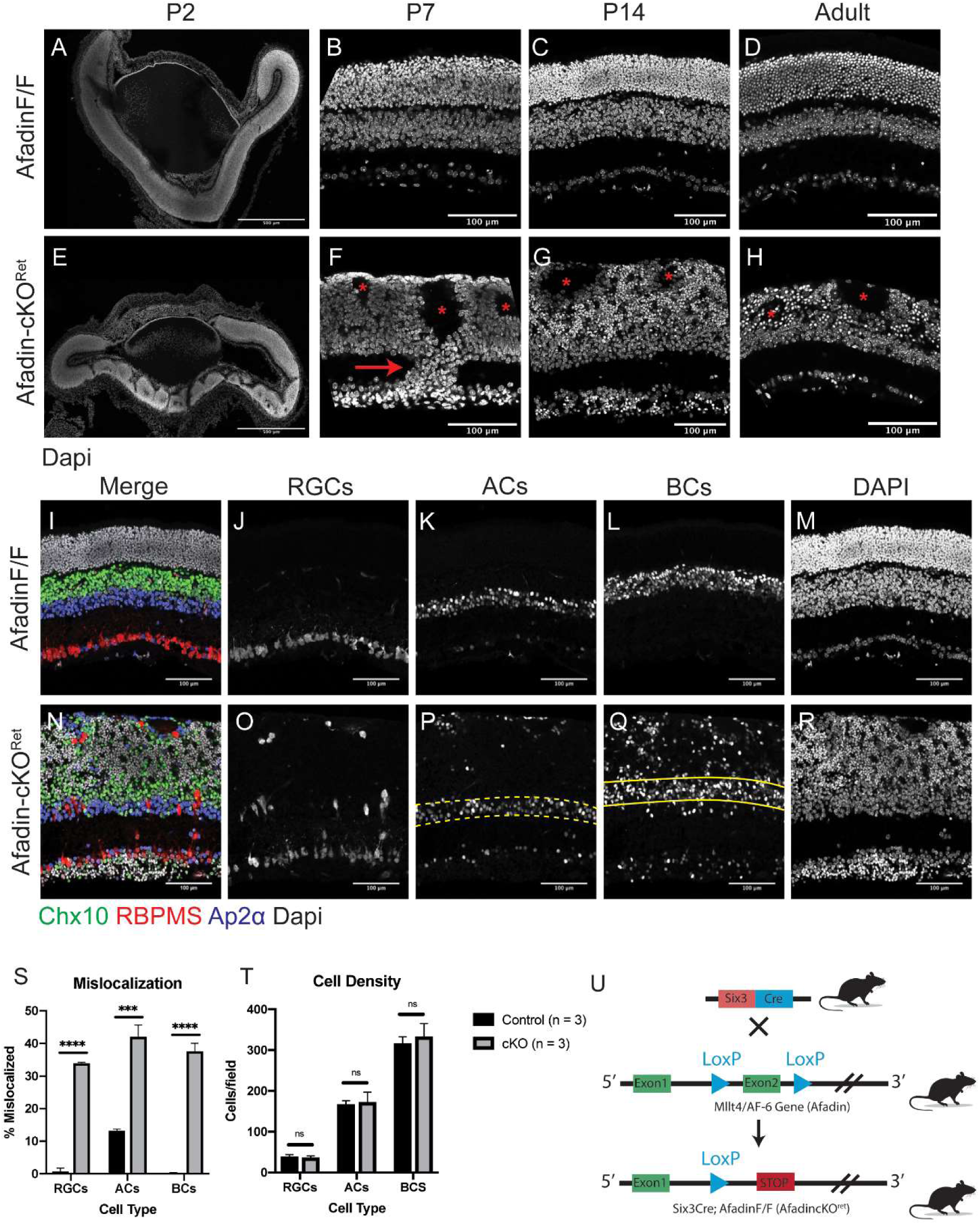
Retina-specific Afadin conditional mutants disrupt the cellular layer organization. **A-H,** Postnanatal time-course of mouse retinal cryosections in Afadin control (Afadin^F/F^) and Afadin knockout (Afadin-cKO^Ret^). The Six3-Cre-Afadin knockout (**E-H**) displays disruption of the tri-neuronal layer organization. As a result, it leads to a fused singular outer layer (fused INL/ONL) compared to control retinae (**A-D**), which contain three distinct nuclear laminae separated by two plexiform layers. Rosettes (**F-H, asterisks**) in the fused INL/ONL are visible from as early as P2 to adulthood and are devoid of cell bodies, and primarily contain neurites (see Figure 2G, 2K, 2L) (**F-H**). The IPL is retained in Afadin-cKO^Ret^ but contains columnar-like structures of displaced neurons (F, arrow). In adult Afadin-cKO^Ret^ mice (**H**), there is significant shrinkage of the fused INL/ONL (see Figure 4E-H). Scale bars (**A-H**): 100μm. **I-T,** Afadin conditional knockout results in the mis-localization of major retinal cell types. In cross-section view (**I-R**), the control retina displays stereotypical lamination of three major cell types: bipolar cells (Chx10), retinal ganglion cells (RBPMS), and amacrine cells (AP2a) (**I-M**). In control retinae, RGCs and BCs stayed in the GCL and INL strictly, with very little displacement. ACs have about 13.2±0.4% displacement (S). In contrast, Afadin- cKO^Ret^ showed aberrant localization of three major cell types (**N-R**). RGCs, ACs, and BCs display 33.9±0.4%, 42.0±3.7%, and 37.6±2.4%, respectively (**S**). Across three replicates there was no significant difference between cell counts across the three cell types (**T**). Unpaired two-sided student’s t-tests; n.s., not significant; ****, p < 0.0001; ***, p < 0.001. Data presented as mean percentage mislocalized±SEM. Mislocalization and cell density quantification were obtained from P14 mice. n=3 mice in each condition. Scale bars (**I-T**): 100μm. **U.** Generation of Afadin-cKO^Ret^ mice. Six-Cre transgenic mouse crossed with Afadin conditional knockout mouse. Exon 2 is flanked by LoxP sites, enabling Cre-mediated deletion, resulting in a frameshift and premature stop codon.

**Figure 2.**
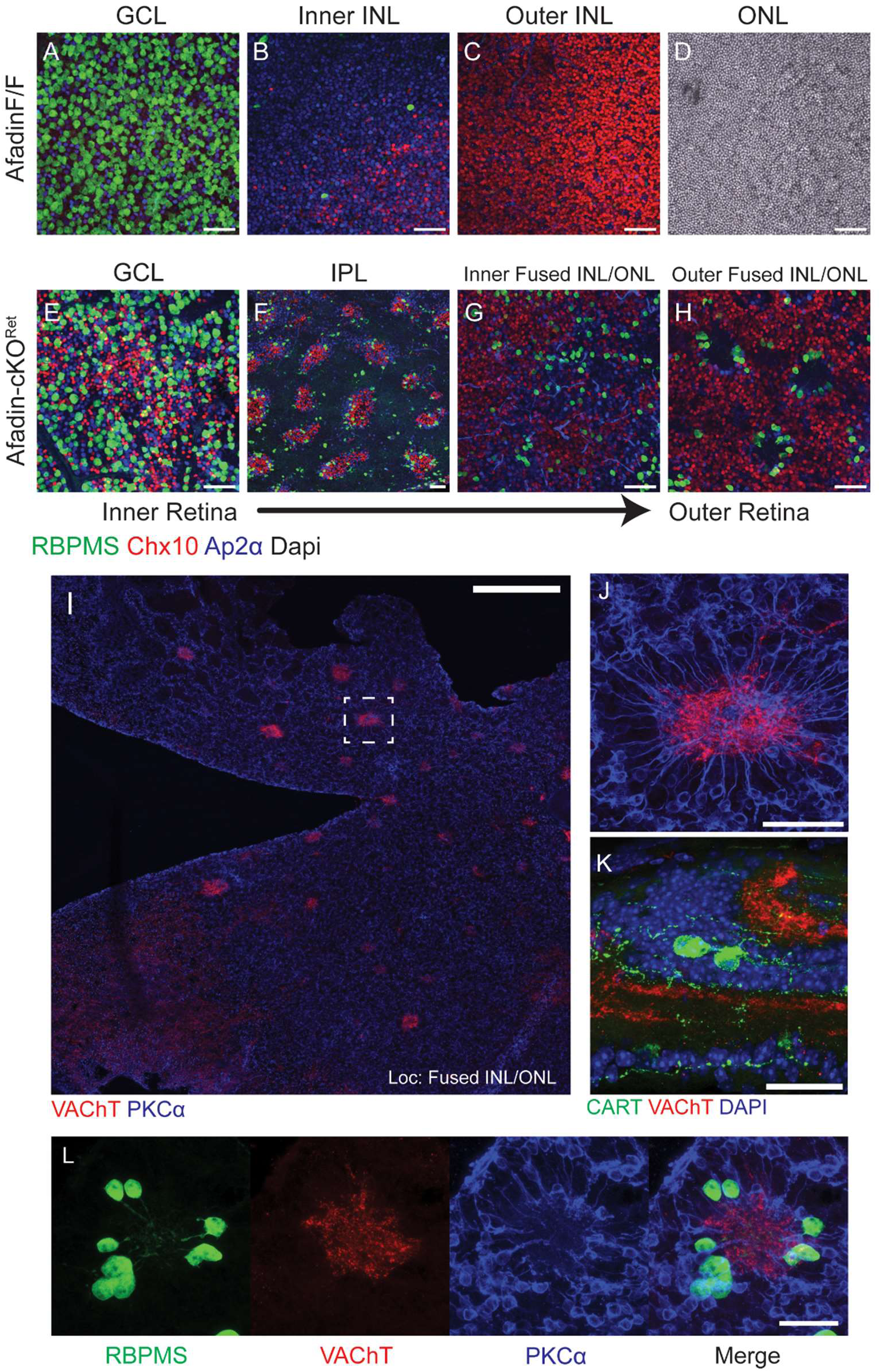
Synaptic rosettes persist despite lateral displacement of cell types. **A-H,** Wholemount retinal cross-sections display lateral displacement of major cell types in Afadin-cKO^Ret^. In a wholemount section view, Afadin-cKO^Ret^ (**E-H**) displays lateral displacement of cell types within unexpected laminae, which are absent in control (**A-D**). Notably, BCs (Chx10) are seen in the GCL (**E**), and RGCs are visible in the fused INL/ONL (**G and H**). ACs are found throughout all laminae in Afadin-cKO^Ret^. The IPL in the Afadin-cKO^Ret^ contains regularly interspaced clusters of RGCs, BCs, and ACs, which form vertical bridging columns (**F**); IPL for the control retina is not shown. The rosettes are visible, with the neurites of RBPMS+ RGCs projecting inward toward the rosette center (**H**). Scale bars (**A-H**): 50μm. **I-L,** Rosettes in the fused INL/ONL have characteristics of an ectopic IPL. Wholemount section (**I**) of an outer region of the fused INL/ONL showing starburst amacrine cell (SAC) processes labeled by VAChT forming a central rosette structure upon which BC processes colocalize. In a 60x magnification of the dashed region in I, the spoke-like processes of rod bipolar cells stained with PKCα are visible (**J**). RGC dendrites also co-cluster in the rosette structure projecting centrally (**L**, RBPMS). A subset of these RGCs are Cartpt-positive which labels ooDSGCs, indicating an ectopic IPL circuit composed of DSGCs is retained at the histological level (**K**). Scale bar (**I**): 300μm; Scale bar (**J**): 50μm; Scale bars (**K-L**): 30μm.

### Mis-localized neurons in Afadin-cKO^Ret^ form an ectopic inner plexiform layer

In addition to scrambled neuronal locations in the wrong cellular layers, we also observed unusual outer layer “rosette” structures and IPL bridging columns in Afadin-cKO^Ret^ that were evident as early as P2. To better characterize these substructures in our model, we utilized wholemount retina sections to obtain an *en-face* view of the IPL and fused INL/ONL (Figures 2E-H). Interestingly, these rosettes retained some degree of canonical neuronal patterning (Figure S1A-F). They comprised radially arranged rod BCs projecting inwards towards ACs and RGCs (Figure 2I-J). The subtype identities of RGCs within the clusters were diverse, including Osteopontin (Spp1)^+^ αRGCs^28,29^, Melanopsin (Opn4)^+^ intrinsically-photosensitive RGCs (ipRGCs)^30,31^, and Cartpt^+^ ON-OFF direction-selective ganglion cells (ooDSGCs)^32^. In addition, dendrites of Carpt^+^ ooDSGCs are projected centrally. They colocalized with starburst amacrine cell (SAC) dendrites (Figures 2J, 2K, and S1D), reminiscent of traditional synaptic pairings at the IPL among these cell types^33,34^. We next quantified the number of “rosettes,” which numbered approximately 160 ± 13 (Figures S2A and S2B) across four Afadin-cKO^Ret^ retinas in adults. This suggested that rosette formation was likely not a random occurrence but was likely a result of compensatory mechanisms of retinal lamination independent of Afadin and its CAM partners.

### Axon pathfinding to central visual targets remains intact in Afadin-cKO^Ret^

Prior work exploring the role of Afadin and N-cadherin in the mouse dorsal telencephalon revealed that loss of either led to severe defects in axonal pathfinding, in addition to neuronal mislocalization and increased progenitor cell proliferation^19,20,35^. We asked whether loss of Afadin in the retina would affect RGC projections to the superior colliculus (SC), one of the primary retinorecipient areas in the mouse receiving input from more than 85% of RGCs^36–38^. To delineate the contributions from the right and left eyes, we administered intravitreal injections with CTB-488 and CTB-555 to label projection neurons, i.e., RGCs from the eyes: these dyes are subsequently transported via RGC axons to the SC (Figure 3A).

**Figure 3.**
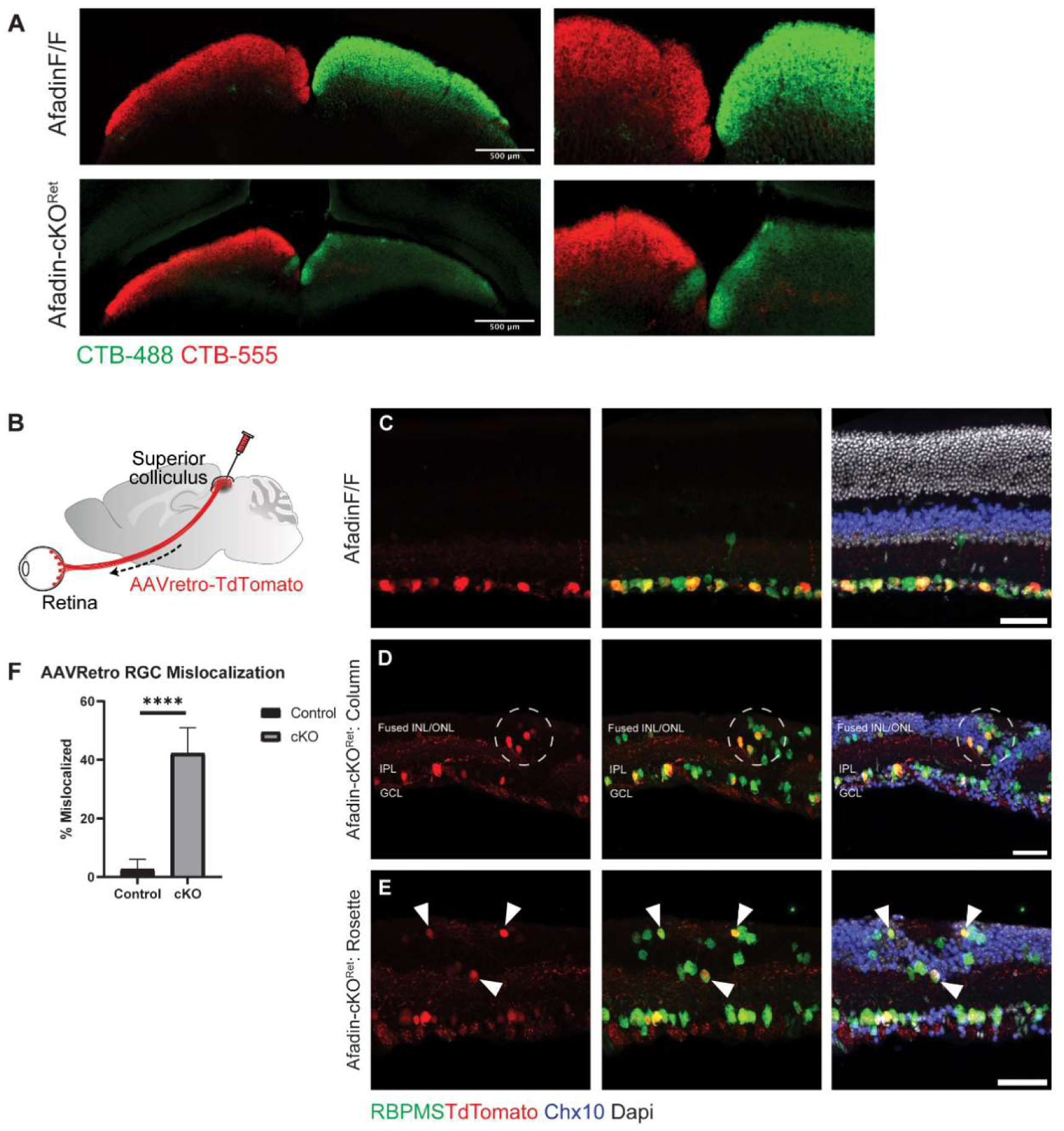
A Significant fraction of displaced RGCs project to the central targets in the SC. **A,** Superior Colliculus (SC) sections labeled with bilateral retina injections. CTB-488 and CTB-555 dye were injected into the left and right eye, respectively, of both AfadinF/F and Afadin-cKO^ret^ mice. An ectopic CTB-488 patch was found close to the midline of the left SC. Scale bar: 500um **B,** Illustration showing stereotaxic protocol. AAV (Retro)-TdTomato was injected unilaterally into the right superior colliculus (SC) of adult mice. The retrograde virus was uptaken by RGC terminals in the SC and selectively labeled RGC somata in the retina. **C-E,** Displaced RGCs in Afadin-cKO^Ret^ send axons to the SC. Retinal cross-sections of Afadin^F/F^ mice show retrograde-AAV labeled RGCs restricted to the GCL layer (**C**). In Afadin-cKO^Ret^ sections, RGCs co-labeled by RBPMS and TdTomato are mislocalized into the fused INL/ONL (dashed circle) and are close to an IPL column (**D**). TdTomato-positive RGCs (arrows) are also mislocalized to a rosette structure in the fused INL/ONL (**E**). Scale bar: 50um **F,** Quantifications of AAVretro-labeled RGC somata. 2.7 ±2.4% of RGCs co-labeled with RBPMS and anti-TdTomato in control mice displayed mislocalization beyond GCL, versus 42.2±8.8% of RGCs in Afadin-cKO^Ret^. Unpaired two-sided student’s t-tests; ****, p < 0.0001.

Interestingly, though RGCs did project to the SC, ipsilateral/contralateral segregation was disrupted. While most (>97%*) projecting axons should cross at the midline in mice and project contralaterally, we noted an unusually high number of aberrant ipsilateral projections in Afadin-cKO^Ret^ (Figure 3A). Using an AAV(Retro)-mediated axonal projection- based retrograde labeling for RGC labeling, we delivered AAV-Retro TdTomato into the brain targets. We observed retrogradely labeled RGC distributions inside the retina (Figure 3B). To our surprise, the retrograde labeling also labeled RGCs displaced into the fused INL/ONL (Figures 3C-E). Approximately 42.2±8.8% of the TdTomato labeled RGCs were mis-localized in Afadin-cKO^Ret^ (Figures 3F), while only a few RGCs are displaced into the INL in the control conditions (Figures 3C and 3F). Altogether, these results suggest that Afadin loss leads to axonal pathfinding deficits; on the other hand, the scrambled RGCs still grow their axons onto the central targets, including the SC.

### Photoreceptor loss in Afadin-cKO^Ret^ disrupts visual function

Associated with Afadin mutants in the inner retina, we inquired about the functional changes associated with such drastic anatomical changes. We observed a progressive photoreceptor loss within the same mutants. Thus, the mutants in their current form prevented us from further inquiring about functional changes associated with the inner retina physiological functions and instead, encouraged examination of how Afadin loss affected photoreceptor lamination and survival. In adult mice at P60 (Figures 4A-B), we observed a close-to-complete loss of central photoreceptors; there was noted peripheral sparing, as Six3-Cre drives primarily central retinal neuroepithelium during development. Using Recoverin, which primarily labels rods and a small subset of BCs, we observed a significant loss of Recoverin^+^ rods in Afadin-cKO^Ret^ mice (Figures 4C-J). The mechanisms leading to the loss of photoreceptors are currently unknown; however, the loss is likely due to the postnatal disruption of the organization and connectivity. Yet, the same mutants caused no major losses of BCs, ACs, and RGCs (Figure 1T). Using the same set of mutants at P60, we sought to obtain a comprehensive measurement of visual function in Afadin-cKO^Ret^ mice. Given the degree of disorganization and photoceptor loss, we noted significantly reduced scotopic and photopic ERG responses (Figures 4M), with flattened a and b waves in scotopic conditions (Figures 4K and 4L). Altogether, these results indicate that loss of Afadin affects photoreceptor stability, leading to significant visual deficits.

**Figure 4.**
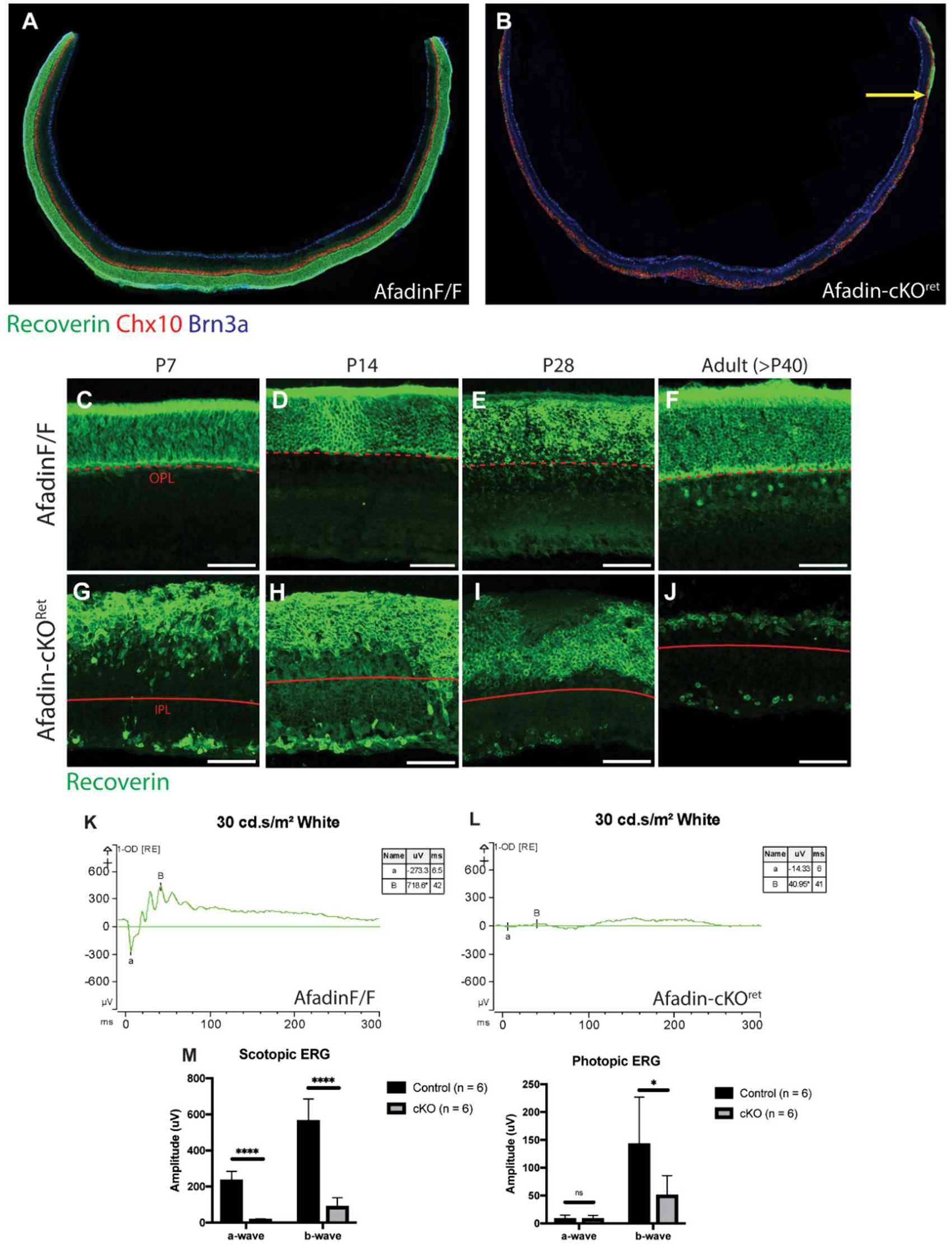
Afadin Mutants lose photoreceptor-mediated visual function in adults. **A-B,** Retinal cryosection displaying thinning and rod photoreceptor loss. Retinal cryosection taken from the central retina of Afadin-cKO^Ret^ (**B**) displays loss of recoverin-positive rod photoreceptors; the photoreceptor layer is maintained in control (**A**). Yellow arrow: displays remnant recoverin-positive patch in the peripheral retina. Scale bars: 500um **C-J,** Postnanatal timecourse of Afadin-cKO^Ret^ displaying progressive photoreceptor loss. Recoverin-positive photoreceptors display aberrant lamination in Afadin-cKO^Ret^ (**G-J**) but not in Afadin^F/F^ (**C-F**). By early adulthood, Afadin-cKO^Ret^ have a near complete loss of photoreceptors (**J**). Dashed red lines indicate the transition from ONL to OPL. Solid red lines indicate the transition from fused INL/ONL to IPL. Scale bars: 50um **K-M,** Representative ERG traces shown for one right eye of both Afadin^F/F^ (**K**) and Afadin- cKO^Ret^ (**L**) mouse when shown a dark-adapted intensity program of 30 cd.s/m^2^ white stimulus. In the control mouse, the a-wave amplitude was -273.3uV, and the b-wave amplitude was 718.6uV. In the mutant, the a-wave amplitude was -14.33uV, and the b-wave amplitude was 40.95uV. The ERG responses were quantified in (**M**). Average a-wave and b-wave responses for Afadin^F/F^ and Afadin-cKO^Ret^ mice after dark-adapting overnight. Under scotopic conditions using 30cd·s/m^2^ flash of white light, the average a-wave response was 235.6 ± 49.0uV for control and 17.0± 4.3uV for Afadin-cKO^Ret^ mice; the average b-wave response was 564.4 ± 121.2uV for control and 89.4 ± 48.3uV for Afadin- cKO^Ret^ mice. Under photopic conditions using 10cd·s/m^2^ flash of white light, the average a-wave response was 8.2 ± 6.7uV for control and 8.3 ± 6.3uV for Afadin-cKO^Ret^ mice; the average b-wave response was 142.7 ± 84.2uV for control and 50.4 ± 35.4uV for Afadin- cKO^Ret^ mice. Unpaired two-sided student’s t-tests; ns, not significant; ****, p < 0.0001; *, p < 0.1. Data presented as mean wave response (uV) +/- SEM between right and left eyes across 6 different mice. ERG quantification was obtained from adult mice. n=6 mice per condition.

## DISCUSSION

Herein, we demonstrate that developmental loss of Afadin significantly affects normal retinal development. We observed profound disorganization of retinal neuronal layers, including RGCs, ACs, and BCs. Interestingly, we also noted the appearance of rosette-like structures containing SACs and radially directed RGC dendrites, suggesting some retention of cellular organization and perhaps indicative of other compensatory mechanisms that regulate synaptogenesis. This disorganization also affects RGC axonal projections onto the superior colliculus. The roles of Afadin in CNS development were examined in other regions of the CNS: In the hippocampus, Afadin loss leads to mislocalization of CA1 and CA3 pyramidal cells; additionally, the spine density of CA1 pyramidal cells neurons is reduced, with the authors postulating that that may be a consequence of reduced cadherin puncta density^5,39^. Similarly, loss of Afadin in the dorsal telencephalon leads to both the dispersion of neural progenitor cells and an increase in their total numbers. Neuronal differentiation was not significantly affected, but cells were localized to inappropriate cortical layers^35^. Interestingly, while we also noted retinal neuron mislocalization into the wrong layers, we did not observe major differences in cell numbers among the retinal subtypes we profiled. This may speak to the varied roles of Afadin or their constituent adherent complexes in different parts of the brain and spinal cord.

It is being increasingly appreciated that cell adhesion complexes play a role in mediating neuronal migration, cellular layer sorting, and synaptogenesis^40,41^. Our work utilized the developing retina to elucidate some of the partners that mediate these interactions.

However, our current study raises several questions that remain to be explored: What other mechanisms drive synaptogenesis within the retina; additionally, what is the role of cell adhesion complexes in regulating axonal pathfinding? The rosette-like structures that retain elements of normal synaptic pairings suggest that synaptogenesis may be a layered process, largely driven by cell adhesion complexes but possibly fine-tuned by other extracellular or intracellular mechanisms, irrespective of activity. Indeed, considerable work has shown that gap junctions and the electrical synapses they facilitate are precursors to the eventual formation of chemical synapses between neuronal pairs. Moreover, through what mechanisms does the loss of Afadin disrupt RGC projections and pathfinding? In the spinal neuroepithelium, deletion of Afadin leads to miswiring of motor circuits, such that certain, typically ipsilaterally projecting neurons instead project bilaterally in the Afadin knockout, leading to loss of left-right limb^42,43^ They propose that this is due to a duplication of the central canal that alters midline signaling within the spinal cord. Whether similar structural abnormalities exist in the superior colliculus and to what extent midline signaling is compromised remains to be examined.

## METHODS AND MATERIALS

### Reagent and Resource Sharing

Requests for reagents and further inquiries may be directed to the corresponding author, Xin Duan.

### AfadincKO ^Ret^ Mutants

All animal experiments were approved by the Institutional Animal Care (IACUC) at the University of California at San Francisco (UCSF). Mice were maintained under regular housing conditions with standard access to food and drink in a pathogen-free facility. Male and female mice were used in roughly equal numbers; no sexual dimorphisms were observed. Animals with noticeable health problems or abnormalities were not used. All ages and numbers were documented. PCR of tail biopsy PCR determined genotypes.

The following mouse lines were used:

1. Six3-Cre expresses Cre recombinase in all of the retina except its far periphery as previously described (Duan 2018 and Lefebvre 2015)^10,15^.
2. Afadin^F/F^ mice were generated by targeting the second exon of Afadin with flanking loxP sites as previously characterized in (Beaudoin 2012)^5^.

### Electroretinogram (ERG) Recording

Six AfadinF/F and six AfadincKO^ret^ mice were dark-adapted overnight, and ERG data was collected in dim red light. Mice were first anesthetized with a combination of ketamine/xylazine/acepromazine (70/10/2 mg/kg), and proparacaine eye drops were administered as local anesthesia. Pupils were then dilated with 1% Tropicamide. The mouse was placed on a heating pad (39 °C) under a dim red light provided by the overhead lamp of the Diagnosys Celeris ERG apparatus (Diagnosys LLC). The light-guide electrodes were placed onto the corneas. For scotopic conditions, we used a white stimulus with 30 cd·s/m^2^ luminance intensity. Signals were captured for 300ms after each step to assess scotopic a- and b-wave function. Following the dark-adapted protocol, we used a photopic intensity ramp protocol to assess function in a light-adapted state. For photopic conditions, we used a white stimulus with 10 cd·s/m^2^ luminance intensity. Signals were captured for 300ms after each step to assess scotopic a- and b-wave function. Following the recordings, each mouse was placed in its home cage on a heating pad (39 °C) to aid recovery from anesthesia.

### Intravitreal Injection

Intravitreal injection protocol, as previously established in (Zhao et al 2023)^28^. Mice were first anesthetized with a combination of ketamine/xylazine/acepromazine (70/10/2 mg/kg). Then, CTB-488 (Invitrogen, C34775) and CTB-555 (Invitrogen, C34776) dye was injected into the vitreous chamber of the right and left eye, respectively, with a fine glass pipette (Sutter Instrument Company). The toxin was allowed to travel anterograde for two weeks before processing brain tissue.

### Stereotaxic Injection into the superior colliculus for AAV-Retro

Stereotaxic injection protocol as previously established by (Tsai et al 2022)^38^. Mice were anesthetized with continuous 2% isoflurane/Oxygen on a stereotaxic setup (Model 940, David Kopf Instruments). Meloxicam (5mg/kg) was administered IP before the surgery and for 2 consecutive days after the surgery. AAV viruses were loaded into a pulled glass pipette connected with a syringe (Hamilton, 7634-O) by a dual ferrule adaptor (Hamilton, 55750-0). Injection speed and volume were controlled by a Microinjection Syringe pump (WPI, UMP3T-1). AAV (Retrograde, RG) -Cag-tdTomato (Addgene, 59462-AAVrg) was injected into the right and left superior colliculus of P60+ adult Afadin^F/F^ (control) and AfadincKO^Ret^ mice. Coordinates for superior colliculi injection: (3.9-4.2 mm posterior, 0.6-0.7 mm lateral to bregma, and 1.4-1.0 mm below the skull). Volume: 600nL for a saturated SC injection. Three weeks later, the mice were humanely euthanized via transcardial perfusion. Brain and retinas were collected in 4% PFA for immunohistochemical analysis.

### Histology and Image Acquisition

Retina section histology as previously established (Toma et al, 2024)^44^. Retina wholemount protocols were previously described in (Duan, 2015, Duan 2018)^15,29^. A lethal overdose of anesthesia sacrificed the mice. The eyes were dissected and post-fixed with 4% PFA. on ice for 1 h and rinsed with 1x PBS. Retinas were analyzed as cryosections and whole mounts.

For frozen sections, tissues were immersed in 30% sucrose for 2 h, then frozen in OCT before sectioning in a cryostat (20μm). For immunohistochemistry, sections were incubated in PBS with 3% donkey serum and 0.3% Triton X-100 for 1-h blocking, followed by primary antibodies overnight at 4°C. For wholemount retinas, tissues were incubated with blocking buffer (5% normal donkey serum, 0.5% Triton X-100 in 1x PBS) overnight, followed by primary antibodies for 2–4 days at 4°C. Secondary antibodies were applied for 2 h at room temperature. Sections and wholemounts were washed with 1x PBS and mounted using Fluoromount-G™ Mounting Medium, with and without DAPI (Invitrogen). Confocal images were acquired using a Zeiss LSM900 (Carl Zeiss Microscopy).

### Wholemount Sectioning

Retina section histology was described in (N. Tsai, M.R.L., X. D, manuscript in preparation). Eyes were collected and fixed in 4%PFA on ice for 30 minutes, dissected to remove the cornea and lens, and then placed back into fresh 4%PFA on ice for another 30 minutes. The sclera was peeled off from each retina, and four radial cuts were made. Retinas were then placed in 30% sucrose/PBS and kept at 4°C until the retinas equilibrated/sank. Retinas were mounted onto a 0.45μm membrane filter (MF-Millipore, HABG01300) and stretched until flat. The retina and filter paper then underwent two cycles of drying and rewetting and 30% sucrose/PBS before drying for 5 minutes. The membrane filter was then trimmed to match the retina size. The retina and filter paper were then adhered onto a stage made through mounting a block of tissue freezing medium (EMS, 72592) by mounting a block of tissue freezing medium (EMS, 72592), which was formed earlier within an embedding mold (Polysciences) onto a cryostat chuck and sectioning to form a flat stage. The retina was then quickly embedded with a thin layer of tissue-freezing medium. The chuck was placed back onto a block of dry ice to solidify the tissue freezing tissue-freezing medium. The embedded retina block with chuck was then incubated at -80 °C overnight. Following equilibration within the cryostat, the retina was cut flat by maintaining the orientation of the chuck with the cryostat blade while preparing the block and subsequent sectioning. Sections of 12um thickness were collected onto superfrost plus slides (Fisherbrand, 12-550-1).

## QUANTIFICATIONS AND STATISTICAL ANALYSIS

GraphPad Prism 9/10 was used to generate all graphs and complete all statistical analyses. Statistical significance definitions: n.s., not significant; *, p < 0.05; **, p < 0.01; ***, p < 0.001; ****, p < 0.0001. All data are presented as means ± SEM unless stated otherwise.

### Mislocalization and Cell Density Quantification

Mislocalization and cell density images were analyzed using ImageJ software (NIH). In practice, every eighth section was systematically sampled during cryostat preparation, thus ensuring coverage of the entire visual field. The boundaries of the INL-IPL, IPL-GCL, and AC_layer-BC_layer were used as landmarks for mis-localization quantifications. Definitions of mislocalization: RGCs were considered mislocalized if present past the INL-IPL boundary (outer retina) or within the IPL columns; ACs were considered mislocalized if present in the GCL, IPL columns or past the AC_layer-BC_layer (outer retina); BCs were considered mislocalized if present in the GCL, IPL columns, or within the AC_layer. Notably, the mislocalization was very robust and apparent to multiple co-authors. 5 images were analyzed for each mouse (3 mice for control and 3 mice for KO). An unpaired two-sided t-test was used to determine the statistical significance of the mislocalization difference between cell types across control and cKO conditions. Data presented as Mean percentage mislocalized ± SEM.

### AAVRetro Mislocalization

Definitions of mislocalization for AAVRetro: TdTomato-positive RGCs were considered mislocalized if present past the INL-IPL boundary (outer retina) or within the IPL columns. Notably, the mislocalization was very robust and apparent to multiple co-authors. An unpaired two-sided t-test was used to determine the significance between control and cKO conditions. Data presented as Mean percentage mislocalized ± SEM.

### Rosette Count

Rosette counts were obtained manually using Cell Counter in ImageJ across 4 adult Afadin- cKO^ret^ mice ranging in age from P40 to P70. Data presented as Mean rosette count ± SD.

## REAGENTS and RESOURCES

| Primary Antibodies |  |  |
| --- | --- | --- |
| Antibodies | Source | Catalog Number |
| chicken anti-GFP | Abcam | ab13970 |
| rabbit anti-RFP | Rockland | 600-401-379 |
| rabbit anti-RBPMS | Proteintech | 15187-1-AP |
| rabbit anti-Recoverin | Millipore | AB5585 |
| rabbit anti-Cartpt | Phoenix Pharmaceuticals | H-003-62 |
| goat anti-Chx10 | Santa Cruz Biotechnology | sc-21690 |
| goat anti-VACHT | Promega | G4481 |
| goat anti-Chat | Millipore | AB144P |
| mouse anti-Brn3a | Millipore | mab1585 |
| mouse anti-PKCα | Abcam | AB11723 |
| rat anti-tdTomato | Kerafast | EST203 |
| Secondary Antibodies |  |  |
| Antibody | Source | Identifier |
| Alexa Fluor 488 | Invitrogen | - |
| Alexa Fluor 568 | Invitrogen | - |
| Alexa Fluor 633 | Invitrogen | - |
| AAV Vectors |  |  |
| AAVrg-Cag-TdTomato | Addgene | 59462-AAVrg |

## ACKNOWLEDGEMENTS

We thank E. Dang and L. Pena for their assistance in animal care; We thank Y.M. Kuo and S.L. Wang for their technical support. We acknowledge (NEI P30EY002162) for vision core support, RPB unrestricted fund to UCSF-Ophthalmology; from Bright Focus Foundation Glaucoma Research Fellowship to M. Z.; from NIH (R01EY030138) to X.D; Glaucoma Research Foundation (Catalyst for a Cure to A.L.T, Y.H. D.S.W, and X.D.).

---

**Supplemental Figure 1.**
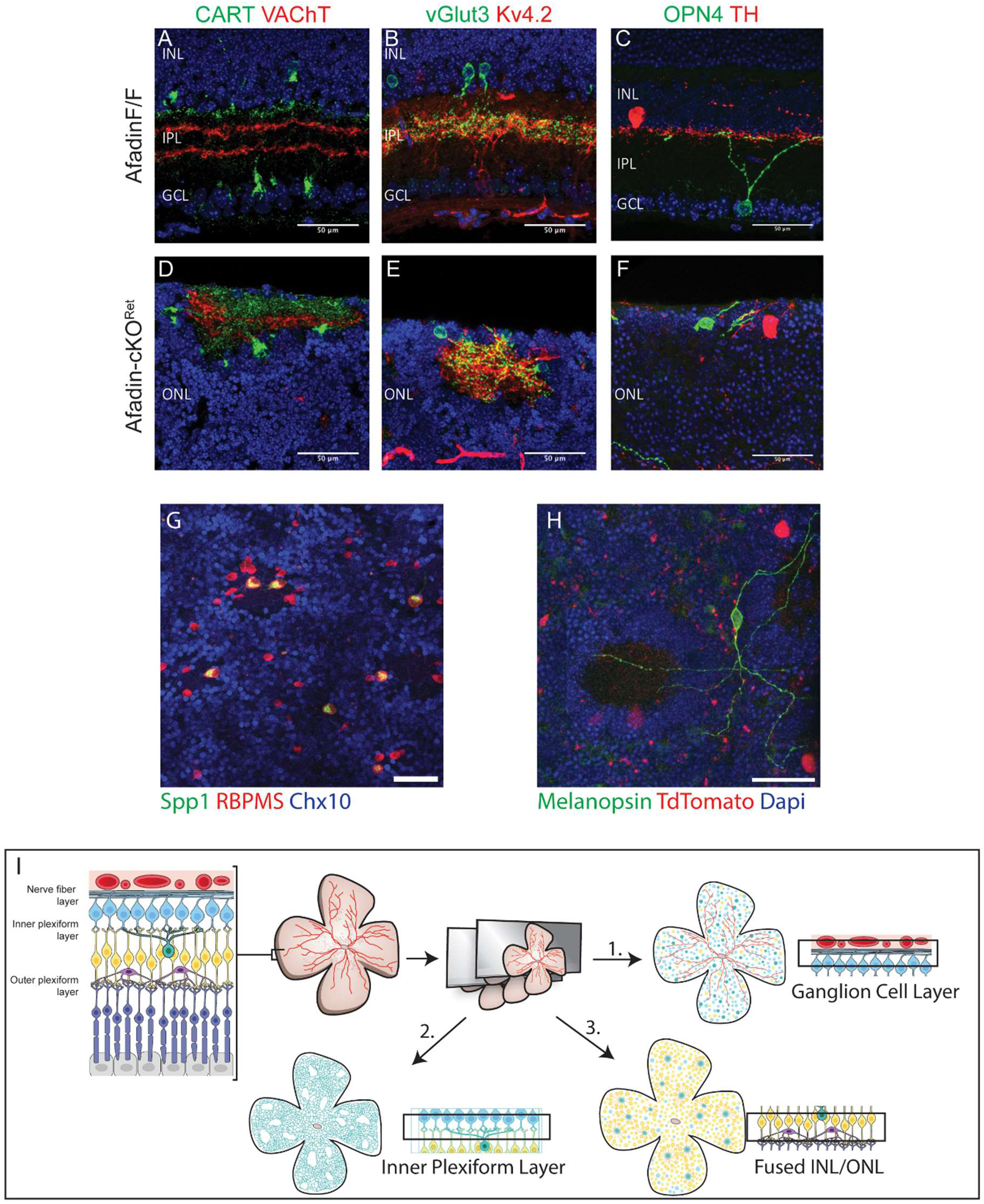
Canonical synaptic pairs persist in Afadin-cKO^Ret^ despite mislocalization. (Related to Figure 2). **A-F,** Canonical amacrine cell (AC) and retinal ganglion cell (RGC) synaptic pairs in the inner plexiform layer (**A-C**) continue to co-fasciculate in Afadin-cKO^ret^ (**S2D-F**), despite ectopic localization of both subtypes in the outer nuclear layer (ONL). These pairings include ON/OFF direction-selective RGCs (Cartpt) and SACs (VAChT) (**left**), glutamatergic ACs (VGlut3) and S3-IPL-targeting RGCs (Kv4.2) (**center**), and dopaminergic ACs (TH) and intrinsically photosensitive RGCs (ipRGCs)(OPN4) (**right**). Scale bars (A-F): 50 μm. **G-H,** In the wholemount view, both αRGCs (Spp1) (**G**) and melanopsin-positive ipRGCs (**H**) were retained near the rosette structures in the fused INL/ONL. Additionally, TdTomato- positive RGCs from AAV-Retrograde injection into the SC are shown (**H**). Scale bars (G-H): 50μm. **I.** A diagram illustrating the major findings and wholemount sectioning procedure is shown (**I**).

**Supplemental Figure 2.**
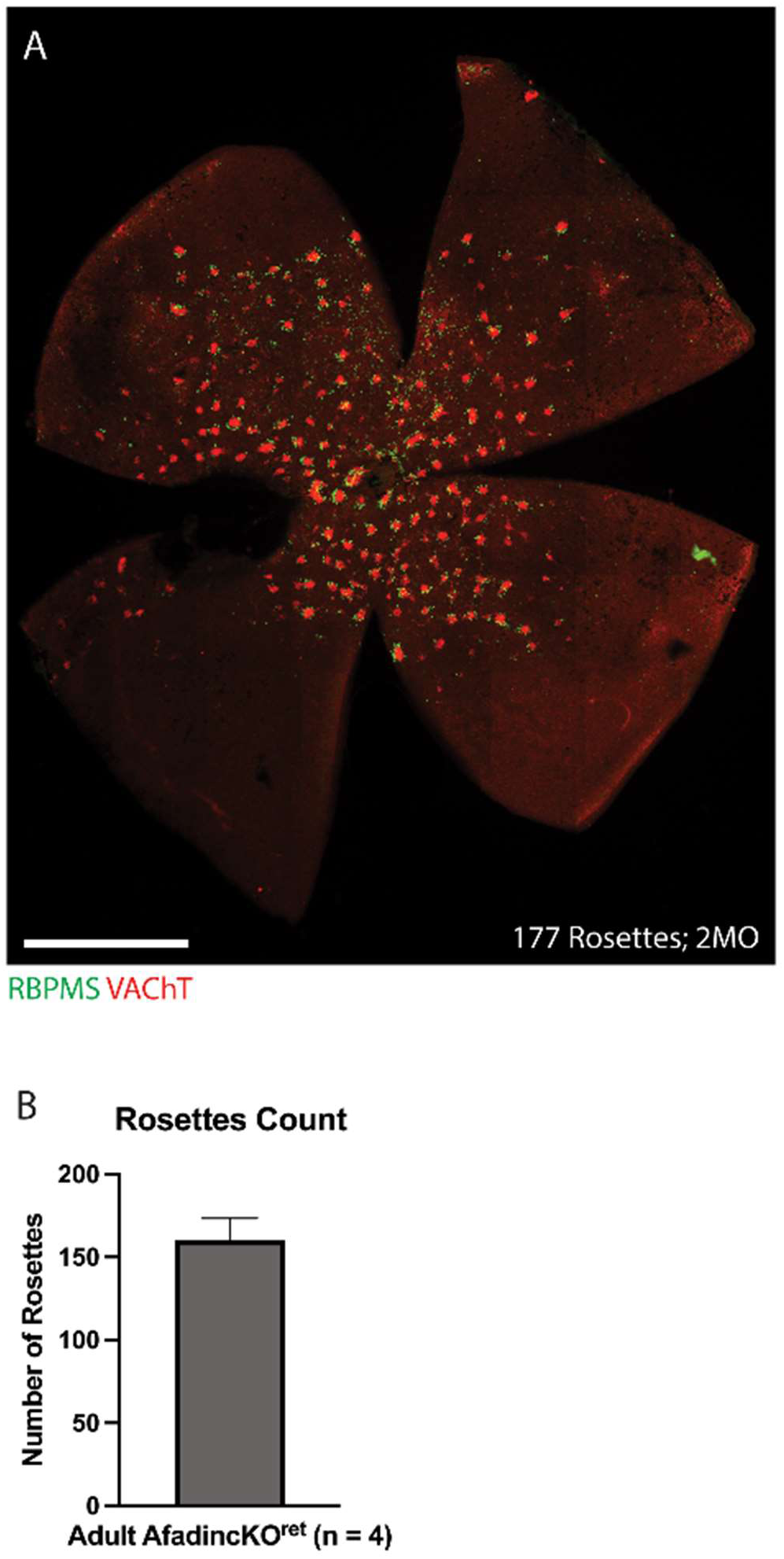
Representative Afadin-cKO^Ret^ Wholemount Representative wholemount displaying rosettes in the fused INL/ONL (**A**): wholemount retina labeled with RBPMS and VAChT aids in the quantification of the number of rosettes per retina. Quantification (**B**) of rosettes across 4 adult Afadin-cKO^Ret^ mice: 160± 13 rosettes across 4 adult Afadin-cKO^Ret^ mice. Data presented as Mean rosette count ± SD.

